# Genome-wide Variants of Eurasian Facial Shape Differentiation and a prospective model of DNA based Face Prediction

**DOI:** 10.1101/062950

**Authors:** Lu Qiao, Yajun Yang, Pengcheng Fu, Sile Hu, Hang Zhou, Shouneng Peng, Jingze Tan, Yan Lu, Haiyi Lou, Dongsheng Lu, Sijie Wu, Jing Guo, Li Jin, Yaqun Guan, Sijia Wang, Shuhua Xu, Kun Tang

## Abstract

It is a long standing question as to which genes define the characteristic facial features among different ethnic groups. In this study, we use Uyghurs, an ancient admixed population to query the genetic bases why Europeans and Han Chinese look different. Facial traits were analyzed based on high-dense 3D facial images; numerous biometric spaces were examined for divergent facial features between European and Han Chinese, ranging from inter-landmark distances to dense shape geometrics. Genome-wide association analyses were conducted on a discovery panel of Uyghurs. Six significant loci were identified four of which, rs1868752, rs118078182, rs60159418 at or near *UBASH3B, COL23A1, PCDH7* and rs17868256 were replicated in independent cohorts of Uyghurs or Southern Han Chinese. A prospective model was also developed to predict 3D faces based on top GWAS signals, and tested in hypothetic forensic scenarios.

## Introduction

The human face plays a pivotal role in daily life. Communication, mutual identification, sexual attraction, etc…all strongly depends on face. It has been long noted that faces bear characteristic features that may surrogate one's ancestry, even in highly admixed populations ^1^. Our recent investigation ^2^ also revealed strong morphological divergence on multiple facial features, including nose, brow ridges, cheeks and jaw, between Europeans and Han Chinese. It is therefore a fundamental and intriguing question to ask: Which genetic variants contribute to the substantial morphological differences among continental populations?

Normal facial shape is known to be highly heritable ^3–5^. However, until recently, very little was known about the genetic basis of common variation of facial morphology. In the last few years, several genome-wide association studies (GWAS) were carried out and multiple face shape associated loci were identified ^6–10^. These studies all based their phenotyping on conventional scalar measurements involving limited number of landmarks. On the other hand, efforts have been paid to use dense 3D face modeling (3dDFM), a novel high-dimensional data format, to fully represent the complex facial shape phenome ^2,11,12^. Peng *et al.* first applied 3dDFM to identify the association between common mouth shape variation and a cleft-lip related genetic locus ^13^. Claes *et al*. showed that numerous genes are associated with complex normal facial shape variation based on 3dDFM ^14^, and proposed the potential of modeling the 3D face based on genotype and its use in forensic practice ^14,15^.

In this study, we aimed at identifying loci on a genome-wide scale that contribute to the divergent facial morphological features between Europeans and Han Chinese. In brief, GWAS were conducted on the polarized face phenotypes along the European-Han dimensions, and Uyghur was used as the study cohort to dissect the genotype-phenotype association. Uyghur is a minority group living in Xinjiang province in China, and was found to have arisen from ancient admixtures between East-Asian and European ancestries at a roughly equal ratio, followed by a long period of isolation ^16–18^. Furthermore, Uyghur facial traits demonstrated a wide range of shape gradients between the characteristic Europeans and Han-Chinese faces ^2^. These properties made Uyghur an ideal group to study the genetic variants of divergent facial features across Eurasia. We performed GWAS in 694 Uyghurs using both landmark based and 3dDFM based phenotypes. Significant loci were replicated in an independent Uyghur sample and a Han Chinese cohort. Next, we carried out a prospective investigation on whether 3D faces could be predicted to certain degree by using the top associated single nucleotide polymorphisms (SNPs). A quantitative model was established to summarize the phenotypic effects of multiple loci and to simulate realistic 3D face models. The prediction model was further tested in hypothetic forensic scenarios to evaluate the potential enhancement of face identification based on DNA.

## Results

The studied cohorts included two independent Uyghur panels (694 and 171 individuals) from Xinjiang China, used as GWAS discovery panel (UIG-D) and replication panel (UIG-R) respectively. UIG-D and UIG-R were genotyped on Illumina Omni ZhongHua-8 and Affymetrix Genome-Wide Human SNP Array 6.0 respectively. In order to obtain a common set of SNPs, the array data of UIG-D and UIG-R was imputed to obtain the whole genome sequencing (WGS) SNPs (see Materials and Methods). The SNPs genotyped on arrays are referred to as genotyped SNPs hereafter, in order to distinguish from the imputed SNPs. In addition, 1504 Han Chinese were sampled from Chenzhou China (HAN-CZ) as a replication panel for candidate SNPs (Table 1). Furthermore, a Han Chinese cohort from Taizhou China (HAN-TZ, 929) and 86 Shanghai residents of European ancestry (EUR) were used as the phenotype reference groups (Table 1). The participants were peer group with 20.02 +/− 2.16 (SD) years old. We collected their three-dimensional facial images and mapped them to a common 32,251 points' spatial dense mesh automatically ^12^. Based on these, we first defined the candidate phenotypes of study. Briefly, the face images were jointly analyzed among EUR, UIG-D and HAN-TZ, and complex face data was decomposed to various phenotype measurements (see Materials and Methods). A candidate phenotype is chosen and termed an ancestry-divergent phenotype if there exists a strong phenotypic divergence between EUR and HAN-TZ, and UIG-D covers a wide range in-between (Fig. 1). Three types of ancestry-divergent phenotypes were defined. First, ten inter-landmark distances were selected (see Materials and Methods; Fig. S1A and Tables S1-S3). The other two types of phenotypes were based on decomposing the high dimensional 3dDFM data. Briefly, we first extracted six facial features, namely, the brow ridge, eyes, side faces, cheeks, nose and mouth, grossly based on the reported high among-population differentiation ^2^ (Fig. S1B). The extracted 3dDFM data was decomposed by either partial least square (PLS) or principle component (PC) analysis ^2^, and the PLS model and PC model that defined the strongest segregation between EUR and HAN-TZ were selected as ancestry-divergent phenotypes, hereafter termed as sPLS and sPC respectively in each feature (see Materials and Methods; Fig. 2, Figures S2-S4 and Table S4). In total, six sPLS and sPC phenotypes were defined, corresponding to the six facial features.

**Figure 1.**
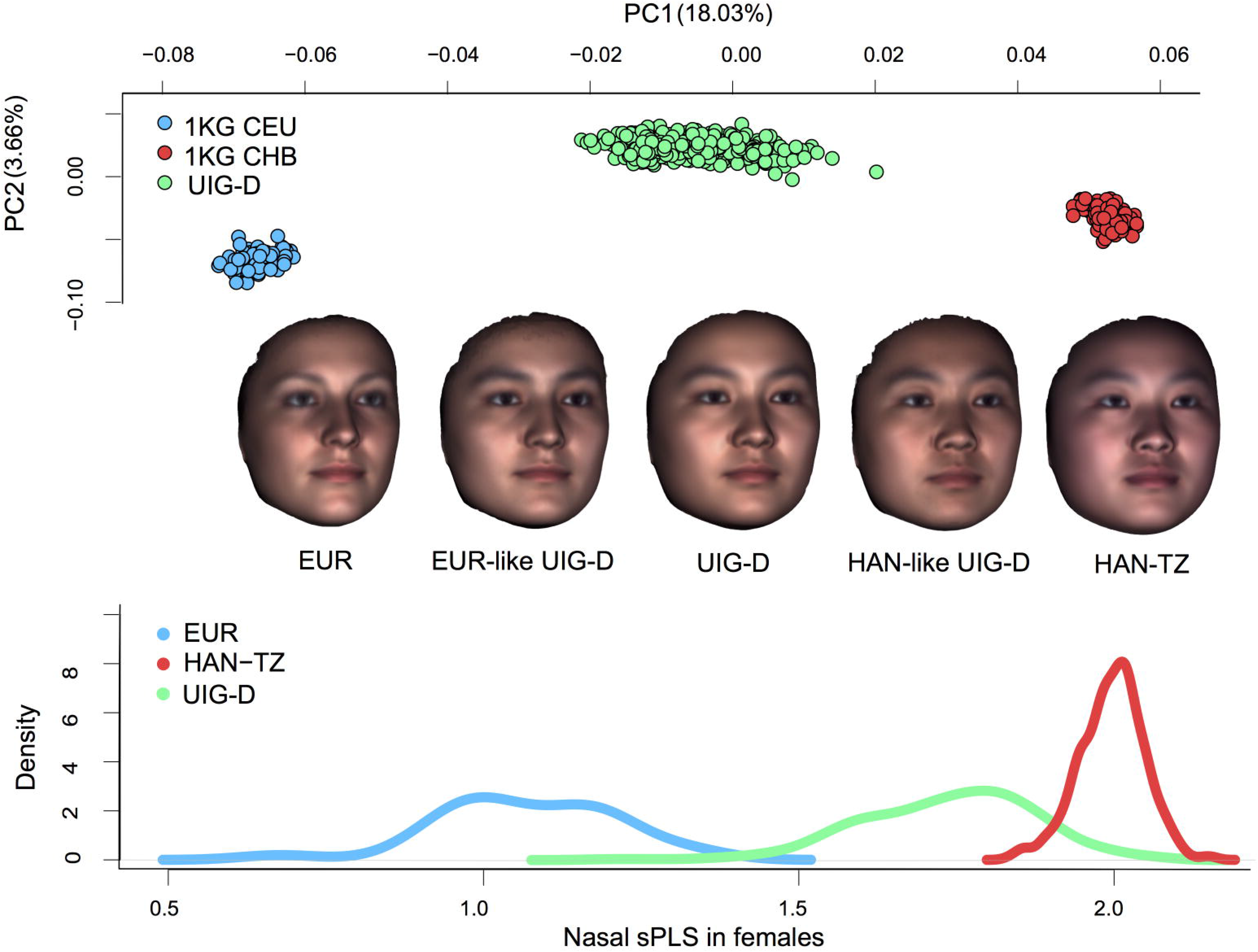
Overall scheme of the study design. In the top panel, the genetic structure of three Eurasian populations was analyzed by PCA based on the 1KG genome data of 97 CHB (red), 85 CEU (blue) and the whole-genome sequencing of 694 UIG-D (green). Clear clustering can be seen based on the ethnic backgrounds. In particular, UIG-D individuals clearly lie in the half way between CEU and CHB along PC1, all consisted of a roughly equal ratio of CHB and CEU ancestries. Compared to the genetic composition, Uyghur individuals exhibit broad gradients of admixture in the facial phenotypes. The middle panel shows the average face models for EUR, European-like Uyghurs (EUR-like UIG-D), UIG-D, Han-like Uyghurs (HAN-like UIG-D) and HAN-TZ from left to right. EUR-like UIG-D and HAN-like UIG-D were obtained by averaging over 20 UIG-D individuals visually accessed to resemble Europeans or Han Chinese. The bottom panel shows the distribution of sPLS in nose, revealing a distinct segregation between EUR and HAN-TZ and a wide spread of UIG-D stretching between EUR and HAN-TZ along this phenotype dimension. In this study, the highly divergent phenotypes as shown above were selected and tested for association loci genome-widely in UIG-D.

**Figure 2.**
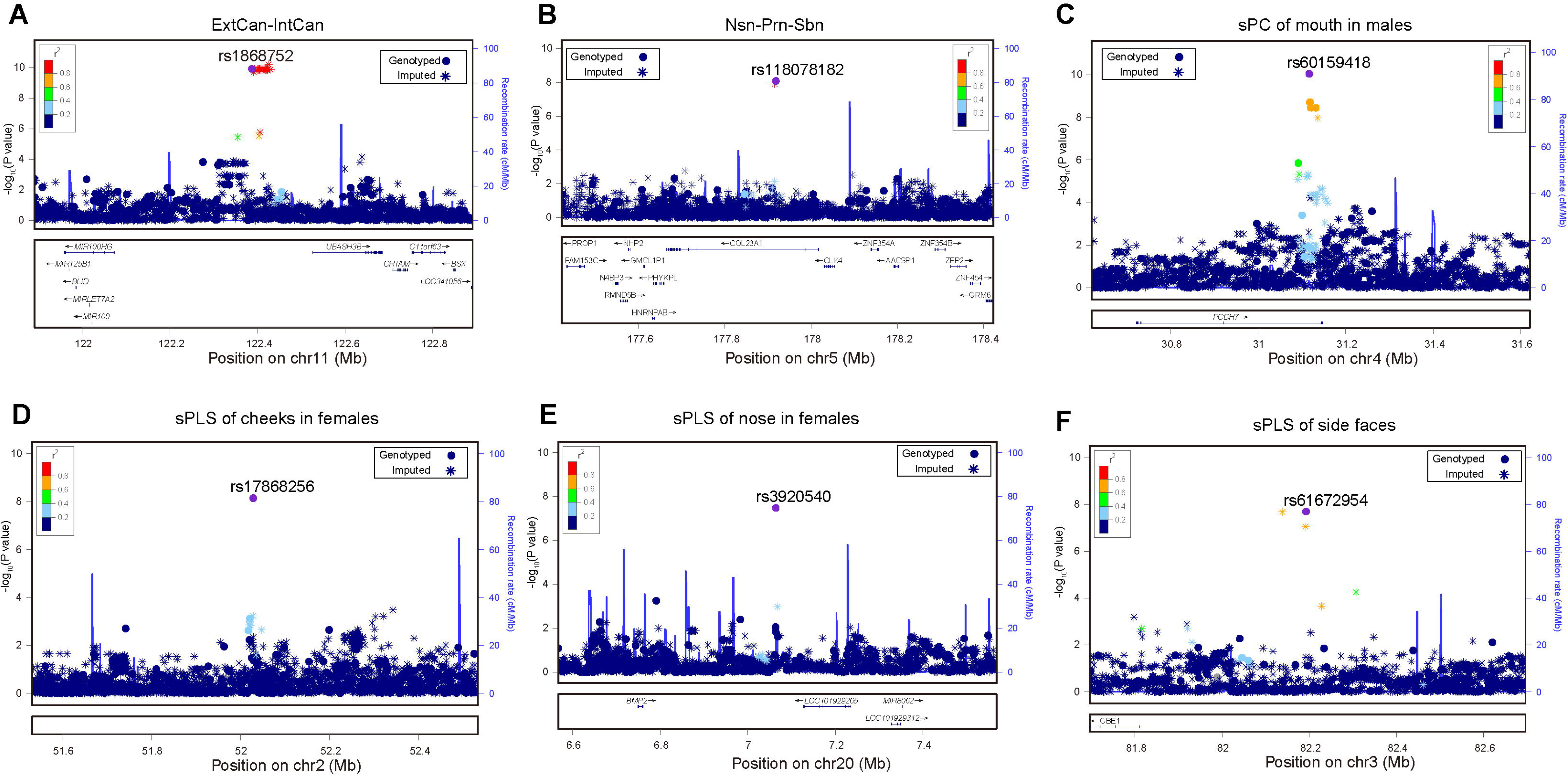
Six genomic regions harboring SNPs of genome-wide significant associations with facial shape. (A) 11q24, (B) 5q35, (C) 4q15, (D) 2q16, (E) 20q12, (F) 3q12. The LocusZoom plots show the association (left y-axis) with corresponding ancestry-divergent traits labelled on the top. Genotyped SNPs are denoted by circles and imputed SNPs are denoted by stars. The plots are given for the 500kb flanking region centered by the genotyped index loci indicated by purple circles. Color gradient of SNPs depict the linkage disequilibrium (r) between each SNP and index SNP.

**Table 1.** Characteristics of study samples.

| Location | Abb. | For | Ethnic | Total N | Male N | Mean Age | Range (s.d.) |
| --- | --- | --- | --- | --- | --- | --- | --- |
| Urumchi, China | UIG-D | discovery | Uyghur | 694 | 270 | 20.09 | 17-25 (1.24) |
| Urumchi, China | UIG-R | replication | Uyghur | 171 | 63 | 20.55 | 17-25 (1.42) |
| ChenZhou, China | HAN-CZ | replication | Han Chinese | 1,504 | 424 | 19.69 | 17-32 (1.67) |
| TaiZhou, China | HAN-TZ | discovery | Han Chinese | 929 | 363 | 19.81 | 17-25 (0.95) |
| Europeans living Shanghai | EUR | discovery | European | 86 | 57 | 27.59 | 16-42 (5.59) |
| N, sample size |  |  |  |  |  |  |  |

### Genome-wide association studies of Eurasian facial shape

We carried out sex-stratified and sex-mixed GWAS in UIG-D on 8,100,752 genome-wide SNPs for the 22 ancestry-divergent phenotypes (see Materials and Methods; Table S5). In general, the lower *P* value inflation assessed by Quantile-Quantile (Q-Q) plot ^19^ analyses revealed little subpopulation stratification (Fig. S5). Six regions revealed signals of genome-wide significance ^20^ at the level of *P*<5 × 10^−8^ (Fig. 2 and Table 2). We focus the discussion on the genotyped SNPs of the highest signal (index SNP) within each region, as their genotypes were more reliable. These include rs1868752 (at 11q24.1) associated with distance between external canthus and internal canthus (ExtCan-IntCan) in mixed genders, rs118078182 (on *COL23A1* at 5q35.3) associated with distance of Nasion point-Pronasale-Subnasale (Nsn-Prn-Sbn) in mixed genders, rs60159418 (on *PCDH7* at 4p15.1) associated with mouth sPC in males, rs17868256 (at 2p16.3) associated with cheek sPLS in females, rs3920540 (near *BMP2* at 20p12.3) associated with nasal sPLS in females, and rs61672954 (at 3p12.2) associated with the sPLS of side-faces in mixed genders. In order to control for potential confounding effects from varying ancestry makeup with mean contribution of ~ 50:50 (CEU:CHB, 0.05 s.d.) (Fig. S6), we inferred the ancestry proportions for each UIG-D individual. The six signals remained after accounting for the inferred ancestry in the association model (Table S6). The genome-wide p-values of the index SNPs should be taken as nominal, as in total 66 GWA analyses (22 traits in females, males and gender mixed) were carried out in total. Noticing that the ancestry-divergent phenotypes were highly correlated among each other (Table S7), we applied Benjamini-Hochberg procedure for FDR ^21^ to address the multiple testing problem (see Materials and Methods). Four index SNPs remained significant genome-widely, including rs1868752, rs118078182, rs60159418 and rs17868256 (Table 2, Table S8).

**Table 2.**
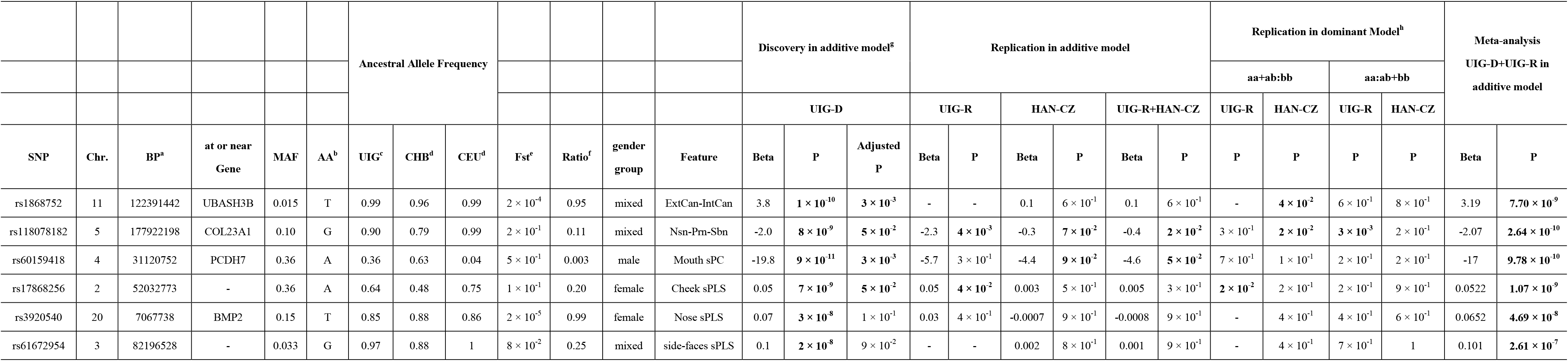
SNPs with GWAS signals and their narrow-sense replications.

a NCBI build 37

b Ancestral allele

c in meta-UIGhurs (UIG-D + UIG-R)

d 1KG phase 3 database

e Fst calculated between CHB and CEU in 1KG phase1 release 2 database

g in additive linear regression model, homozygote ancestral alleles as 0, heterozygote as 1, homozygote derived alleles as 2

h a allele stands for ancestral allele

“-” indicates null results in genes or null test in replications

Ideally, the effects of shape related loci should be directly visualized on facial images. We modeled the effects of index SNPs using heat plots as well as extrapolated faces ^22^ (Fig. 3, Movies S1-S6). As can be seen in Figure 3A (Movie S1), the extrapolated face towards the effect of rs1868752T had narrower eyes (smaller ExtCan-IntCan distance) compared to the G allele, resulting in a substantial displacement on the X axis; G also seemed to be associated with elevated nose ridge. SNP rs 118078182 showed an obvious impact on the nasal shape along the Y and Z axes. Compared to rs118078182A, rs118078182G seemed to make the nose longer and more protrusive (taller) from the face, consistent with the association with Nsn-Prn-Sbn distance (Fig. 3B, Movie S2). For rs60159418 in males (Fig. 3C, Movie S3), the main shape changed the mouth along the Y axis, followed by Z axis. Allele G seemed to make the whole mouth area recessed from the face plane; in comparison, the mouth-chin curve bended convexly from the facial plane in the extrapolated face of the A allele. rs60159418 also seemed to influence other facial features: for G allele, nose and chin looked relatively protrusive outwards, and eye brow ridges seemed to elevate. The SNP rs17868256 mainly affected the shape of cheeks (Fig. 3D, Movie S4), with G allele associated to laterally expanded cheeks, making the face look wider on the X axis. On the Y axis, rs17868256G also seemed to lift the cheek protrusion upwards. For SNP rs3920540 in females (Fig. 3E, Movie S5), G allele was mainly associated with repressed nasal bridge and nasal tip along Z axis compared to T allele; G allele also seemed to link to more protrusive chin on the extrapolated face. The most notable effect of rs61672954 occurred around the jaw lines (Fig. 3F, Movie S6), with the A allele associated with stronger jawlines and therefore comparatively wider lower face than the extrapolated face of G allele.

**Figure 3.**
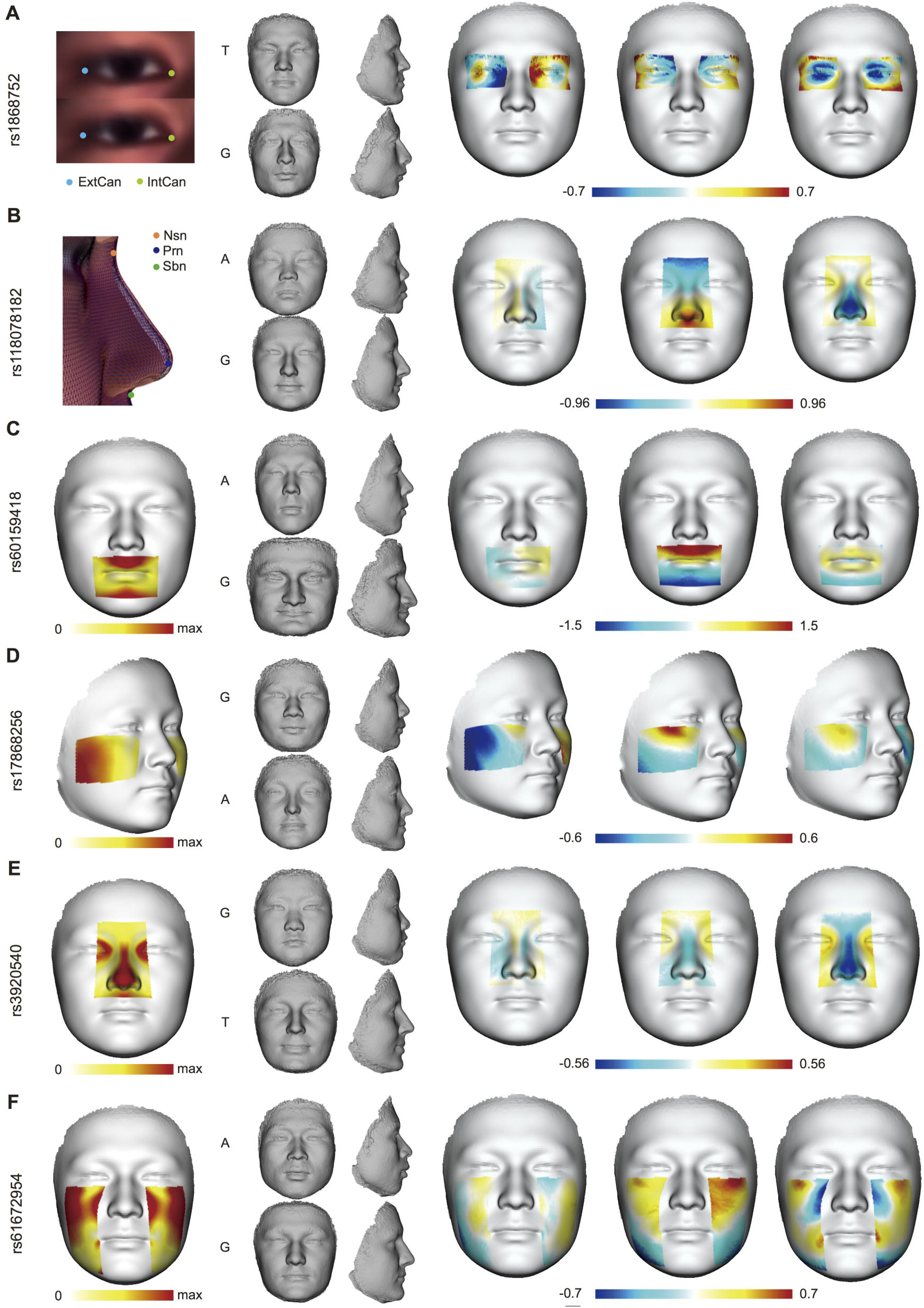
Heat plots and extrapolated faces of six index SNPs affected on responding partial shape. (A) the association of rs1868752 with the distance between ExtCan and IntCan, (B) the association of rs118078182 with the distance of Nsn-Prn-Sbn, (C) the association of rs60159418 with the sPC of mouth in males, (D) the association of rs17868256 with the sPLS of cheeks in females, (E) the association of rs3920540 with nasal sPLS in females, (F) the association of rs61672954 with the sPLS of side faces. For each SNP, the first face shows the general effect on the corresponding feature as the displacement of landmarks or meshes. The mid-panel of four miniature faces gives the extrapolations towards the Han trend on the top, or the European trend on the bottom, with the associated allele labeled at the left side. The extrapolated faces were morphed by exemplifying the difference between the average faces of the opposite homozygotes if both are more than 10% frequent in UIG-D, or the major homozygote and the heterozygote if otherwise. The last three faces depict the signed displacement of the average faces of the fore-mentioned genotypes in X, Y and Z axes; obtained by subtracting the average face of European-trend from that of the Han-trend.

### Replication studies and meta-analysis

We replicated the six GWAS significant loci in an independent Uyghur cohort (UIG-R) and a Han Chinese cohort (HAN-CZ) (Table S9). The former has the same ethnic background with UIG-D ^16–18^ and the latter represents a pool of Han Chinese ancestry from southern China ^23^. Face is highly complex, and the effects of genetic variants on face can be subtle and strongly depend on other factors such as ethnicity and gender ^14,24,25^. On the other hand, face related genetic loci may be pleiotropic, e.g., that a single variant may influence facial morphology on different parts and/or in different ways ^6,7,14,26,27^ In view of this, we defined two types of association replications: the narrow-sense and broad-sense replications. Narrow-sense replication stood for the association signals replicated on exactly the same ancestry-divergent phenotypic measurement; whereas broad-sense replication required the index loci to show evidences of association with any shape changes in the same facial feature. In this study, both the narrow- and broad-sense replications were conditioned in the same gender group as for the discovery panel. As a result (Table 2), the association of rs1868752 with ExtCan-IntCan exhibited narrow-sense replication (*P*=0.04) in a dominant model in HAN-CZ. The SNP rs118078182 showed narrow-sense replications in an additive (*P*=0.004) and a dominant model (*P*=0.003) in UIG-R, in a dominant model (*P*=0.02) in HAN-CZ, as well as in the additive model of UIG-R and HAN-CZ combined (*P*=0.02). For rs60159418, the narrow-sense replication was tested by projecting the UIG-R and HAN-CZ 3dDFM data to the sPC of mouth where the GWAS signal was found in UIG-D males, and revealed marginal significance in HAN-CZ (*P*=0.09) and combined group of UIG-R and HAN-CZ (*P*=0.05). For rs 17868256, the narrow-sense association in females was successfully replicated for cheek sPLS in UIG-R for both the additive model (*P*=0.04) and a dominant model (*P*=0.02). The other two index loci didn't show evidences of narrow-sense replication.

To systematically test the broad-sense replication, we carried out pair-wise shape distance (PSD) permutation as previous proposed ^13^ and PLS-based permutation for the 3dDFM data (see Materials and Methods). Table 3 summarized the results of broad-sense replications. In general, these tests confirmed the results of narrow-sense replications, showing that rs1868752, rs118078182, rs60159418 and rs17868256 affect the overall shapes of the corresponding features. Furthermore, rs3 920540, the nose related locus that failed to replicate in the narrow-sense test, turned out to significantly affect the overall nasal shape in PSD permutation test (*P*=0.01) and PLS-based permutation test (*P*=0.03) in HAN-CZ.

**Table 3.**
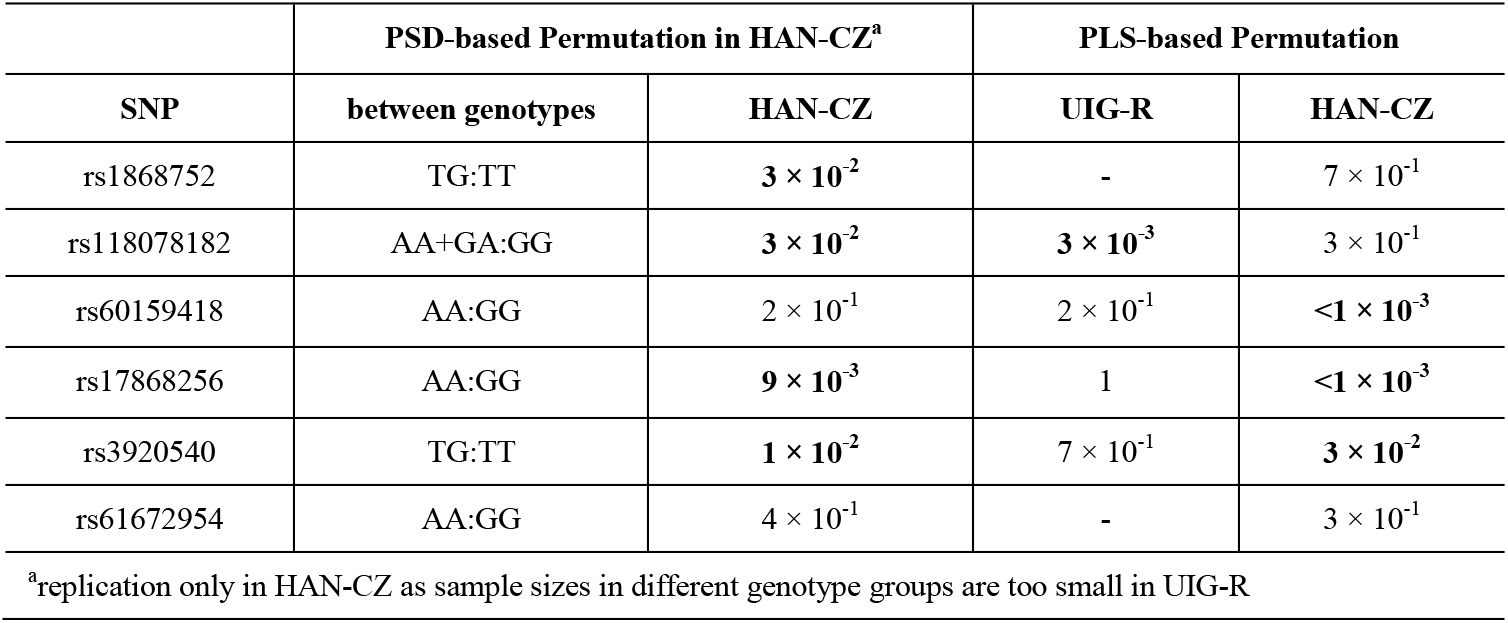
Broad-sense replications of SNPs with GWAS signals.

Visualization in UIG-R and HAN-CZ revealed highly consistent effects of the ancestry-divergent variants as in UIG-D ^22^ (Fig. 4), despite the distinct ethnicity of HAN-CZ. Intriguingly in UIG-R, the effects of some index SNPs were strongly persistent not only within the facial features of GWAS signals, but also across the whole face. In HAN-CZ, similar influence on the whole face could also be observed. In particular, rs1868752T was involved in recessive eye-sockets and repressed nasal bridge in all three cohorts (Fig. 4A); rs118078182A seemed to make the whole face more flat and round in addition to its effects on nasal shape (Fig. 4B); Other than affecting the mouth shape (Fig. 4C), rs60159418G was also associated with stronger brow ridges and more protrusive chin among the three groups. These indicated that the identified association signals were authentic, and the candidate variants were in general highly pleiotropic.

**Figure 4.**
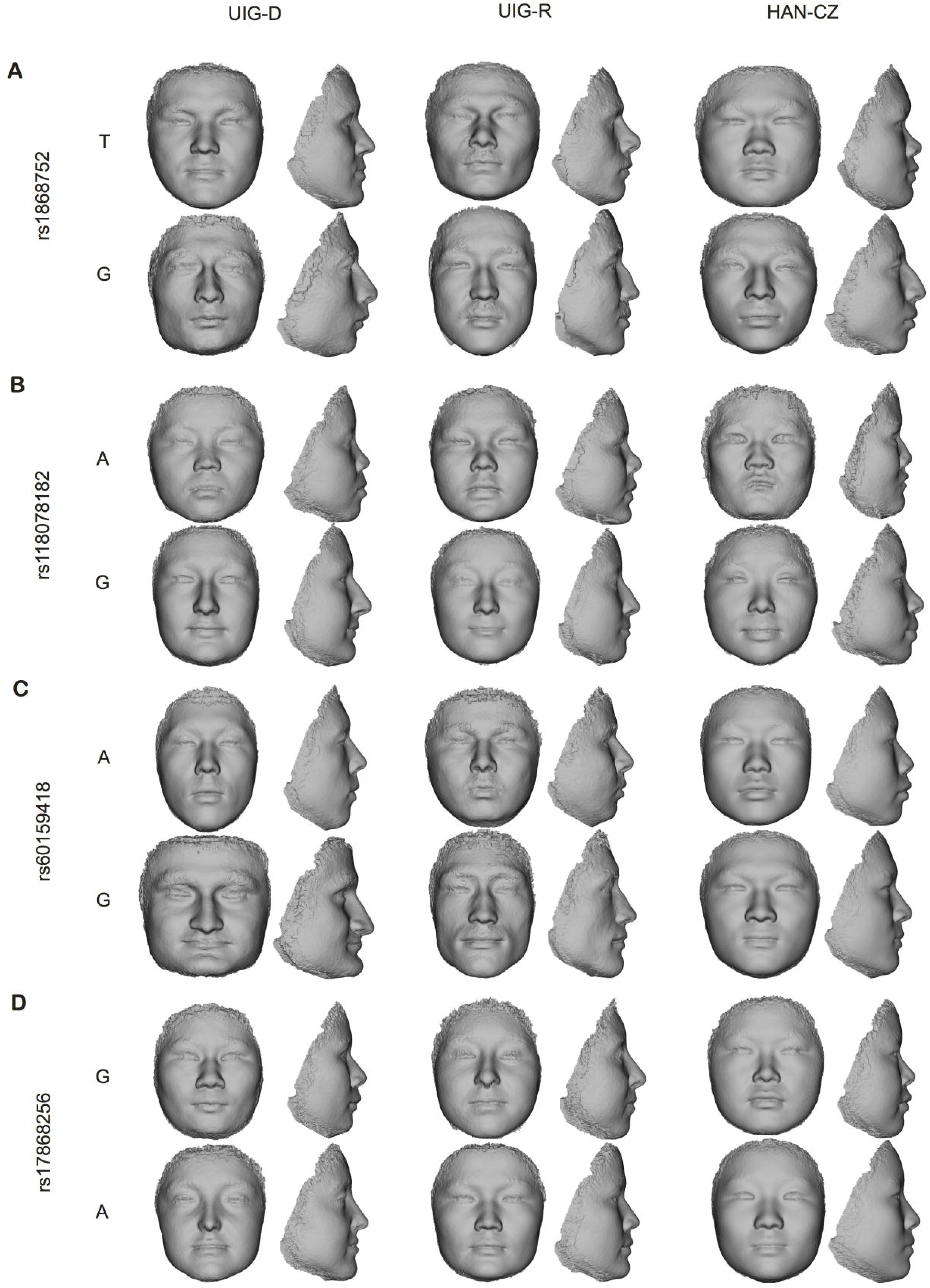
Visualization in UIG-R and HAN-CZ revealed largely consistent effects of the ancestry-divergent variants as in UIG-D. For the four index loci, (A) rs1868752, (B) rs118078182, (C) rs60159418 and (D) rs17868256, we compared the extrapolated faces in UIG-D, UIG-R and HAN-CZ from left to right. For each locus, the top faces in the trend of Han Chinese and the bottom ones are in European trend for the corresponding feature.

### Replication of reported variants of facial variation

We explored whether the candidate SNPs that previously reported to affect normal facial variation ^6−8^ also showed signals of association in our combined Uyghur cohort (UIG-D + UIG-R). Replications were carried out either on the original or related measurements for 12 SNPs (Table S10). Notably, numerous candidate loci were re-validated to varying degrees. Briefly, rs4648379 in *PRDM16,* rs7559271 in *PAX3,* rs2045323 in *DCHS2,* rs17640804 in *GLI3,* rs805722 in *COL17A1* and rs927833 in *PAX1* reported to affect nasal phenotypes in different ethnic groups ^6–8^, were found to also modulate normal nasal shape in Uyghurs (Table S10). The SNPs rs3827760 in *EDAR* and rs6184 in *GHR* that were previously linked to mandibular shape variation turned out to be significant or marginally significant in our study ^8,27,28^ (Table S10). Interestingly, the SNP rs642961 in *IRF6,* previously found to be associated with the mouth shape in Han Chinese females ^13,29^, also showed marginal significance (*P*=0.05) in Uyghur females but not in males or mixed gender group, implying that the dependence of the genetic effect of rs642961 on gender was shared among different populations.

### A prospective3D face prediction model based on genome-wide SNPs

Hypothetically, if a quantitative trait is highly heritable, a proper model featuring major genetic factors should lead to true prediction ^30–32^. For the purpose of face prediction within Uyghurs, GWA analyses should be carried out on common facial variations in all dimensions. Therefore, in addition to the 22 ancestry-divergent phenotypes, we further included GWA analyses on a comprehensive list of phenotypes of various perspectives within Uyghurs, including 26 inter-landmark, 18 PC and 12 PLS phenotypes (see Supplementary Note). The total number of GWA analyses used for face prediction was exceptionally large (234) and a conventional cutoff for GWAS is impractical. We therefore assumed that SNPs reaching a nominal genome-wide threshold of *P* value < 1 × 10^−6^ were enriched for facial shape related loci. All SNPs that satisfy the threshold were combined into a panel of 277 top SNPs (Table S11). Based on these top SNPs, a simple quantitative model was constructed using UIG-D data. Briefly, for each SNP, a residual face was obtained for each genotype by subtracting the genotype average face by the global average. To compose a predicted face (PF), the 277 residual faces were scaled by a global “effect coefficient” *α* (see Supplementary Note; Fig. S7), and then added to the base face (the average face stratified by gender) according to the specific genotypes of the individual of interest. Details of the model construction can be referred to the supplementary materials (Supplementary Note).

The prediction model was applied to Uyghur individuals from the independent replication panel (UIG-R). The PF and actual faces can be visualized but quantitative evaluation of their similarity is difficult with human observers (Fig. 5A, Movie S7). To formally access the resemblance between predicted and actual faces we constructed a robust shape similarity statistic: the shape space angle (SSA). SSA was defined as the angle between two shapes in the 3dDFM data space ^15,33^ (see Supplementary Note). SSA would achieve 0 if the two shapes were the same. A SSA of 90 degree stood for statistical independence, whereas a SSA greater than 90 indicated that two faces deviated from the mean face in reverse directions. Tests of shape similarity were carried out in UIG-R stratified by gender to see whether the differences from the actual face to their corresponding PF was indeed smaller than to random faces. We used two types of random faces. The first was the random-genotype predicted faces (RGF; see Supplementary Note), adopted from a previous study ^15^. Briefly, a RGF was generated in the same way as the PF, except that the genotype set used for prediction was replaced by a random genotype set permuted from the known frequencies of genotypes in the combined UIG cohort. The other was the random-actual faces (RAF) set, which was obtained by randomly sampling the UIG-R actual faces. We constructed two types of tests, one is to compare the SSA distributions (inter-distribution test) between the random predicted faces and the PF faces; the other test the average similarity score (single-score test) between the PF and actual faces, against the null distributions based on multiple iterations using RGF ^15^ or RAF as prediction (see Supplementary Note). Both tests generated similar results (Figs. 5B and 5C), that the females of UIG-R did not show marked differences between the PF and random predictions (the inter-distribution test *P*=0.87 in RGF and *P*=0.73 in RAF; the single-statistic test *P*=0.35 in RGF and *P*=0.34 in RAF). However, in males, the true prediction significantly outperformed the random predictions in all the tests (the inter-distribution test *P*=0.01 in RGF and *P*=0.04 in RAF; the single-statistic test *P*=0.02 in RGF and *P*=0.02 in RAF).

**Figure 5.**
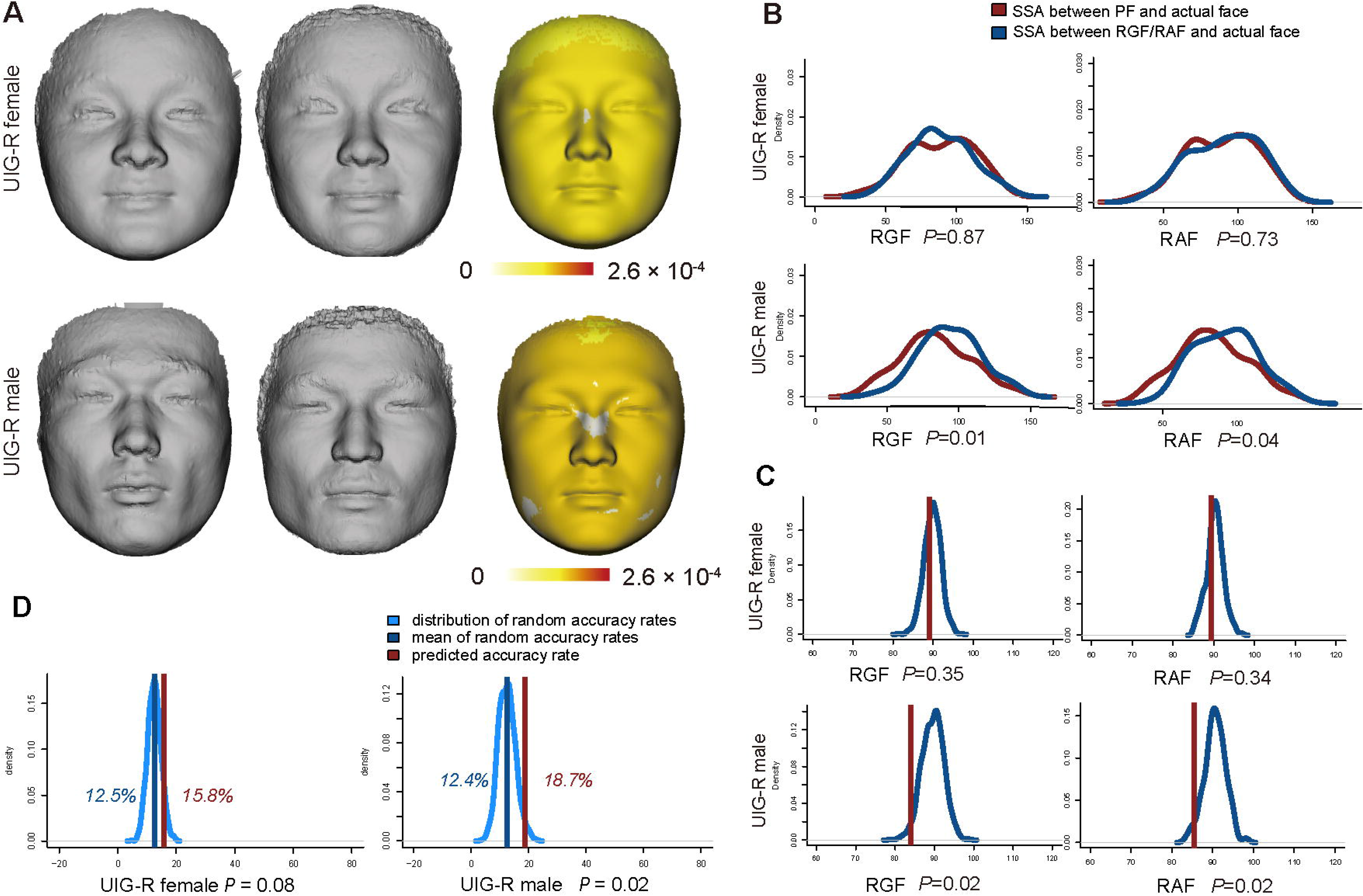
Test of the prediction model in UIG-R. (A) cases of visualization of actual face (left column), the PF (middle column) and the displacement between the pair in heat plot (right column). (B) Evaluation by inter-distance test. The comparison between true SSA distribution (in brown) and random SSA distribution in RGF (left column) or RAF (right column) was tested by Student's Test. (C) Evaluation by single-statistic test. The average SSA determined for the cohorts (in brown) were compared to the RGF (left column) and RAF (right column) distributions under null hypothesis (in dodgerblue). *P* values are the probability of predicted statistic distributed on the relative random normal distribution calculated like normal one-side *P* value. (D) Evaluation of the face prediction in a hypothetic forensic scenario. SSA as accuracy statistics are evaluated in UIG-R females (left column) and UIG-R males (right column). The true accuracy rate based on SSA, in brown, were determined by examining how many cases of successful identification were achieved in all combined iterations. The random accuracy rate was calculated after reshuffling the pairwise corresponds between actual face and PF. This process was repeated 1,000 times to obtain a distribution of random accuracy rates (in dodgerblue) under null hypothesis. The *P* values are the proportion of how many random accuracy rates were larger than true accuracy rate calculated as the empirical p-value.

Intriguingly, face prediction based on genotypes would promote the suspect identification in pragmatic forensic scenarios. We simulated toy forensic scenarios where a single true suspect should be picked up from a group of 8 candidates (Fig. S8). We supposed that the 3D facial data and genotypes could be obtained from each candidate to allow mathematical comparison between the actual and PF. Interestingly, there was a significant increase in accuracy (+6.3%, empirical *P*=0.025) in males compared to the simulated neutral distribution (12.4% ± 0.03SD) assuming no prediction power at all (see Materials and Methods). In females, the prediction did not substantially enhance the accuracy (+3.3%, empirical *P*=0.076) (Fig. 5D).

## Discussion

To our knowledge, this was the first GWAS aimed at identify genetic loci associated with normal facial variations based on complex 3dDFM data that revealed multiple genetic determinants underlying the European-East Asian facial trait divergence. The genome-wide significant loci were located on independent regions and respectively associated with shape of eyes, nose, mouth, cheeks and side faces. We successfully replicated four loci, rs1868752, rs118078182, rs60159418 and rs17868256 in independent cohorts on the same phenotype measurements. In addition, the association signal of rs3920540 was replicated in broad-sense.

SNP rs1868752 is not located in any gene region. The nearest protein coding gene is the ubiquitin associated and SH3 domain containing B (*UBASH3B*) about 140kb distal, which regulates epidermal growth factor receptor (*EGFR*) and platelet-derived growth factor receptor (*PDGFRA*). The longer eye length, consistent to European traits is associated with the derived allele rs1868752G. However, rs1868752G has a low global frequency (< 10%), and the population frequency in CEU is even lower (~1.5%) than in CHB (~4%), suggesting that this SNP does not play a major role in the Eurasian face differentiation. The SNP rs118078182 has the most consistent association signals across different sample panels and test models. This SNP is an intronic variant in collagen type 23 alpha 1 (*COL23A1*). COL23A1 codes a non-abundant trans-membrane collagen, primarily found in head, skin, tendon, and kidney ^34^. A possible role of spatial/temporal regulation in facial morphogenesis was noted for *COL23A1* ^35^. Interestingly the G allele of rs118078182, associated with the European trait of longer and taller nose is almost fixed (99.5%) in CEU compared to the sequentially lower frequencies of ~90% in Uyghur and ~79% in CHB, suggesting that rs118078182 plays a major role in the nasal shape divergence across Eurasia. The other nasal shape related SNP, rs3920540 is approximately 300kb away from the nearest protein coding gene *BMP2,* which is a member of the bone morphogenetic proteins involved in the development of bone and cartilage ^36–39^. The T allele of rs3920540 pertaining to the European trait of taller nose does not seem to differ in frequency (0.86, 0.85 and 0.88 in CEU, UIG and CHB respectively) among the three populations, suggesting that this SNP mainly contribute to the within-group variation of nasal shape. The mouth shape related SNP rs60159418 is situated in an intron of the protocadherin 7 gene (*PCDH7*). *PCDH7* codes an integral membrane protein functioning in cell-cell recognition and adhesion. Previous studies showed that *PCDH7* played a key role in osteoclastogenesis ^40^, and its homologue gene is a pivotal regulator in the head formation of the mouse embryo ^41^. Consistently, the derived G allele that co-occurs with the concaved European mouth shape is almost fixed in CEU (0.96, Table 2), much lower (0.64) in UIG, whereas in CHB the ancestral A allele is the major allele (allele frequency 0.63). These suggest that rs60159418 contributes to the mouth shape differentiation across Eurasia. Notably, rs60159418 is among the most divergent SNPs between CEU and CHB (*Fst_CEU-CHB_* = 0.4), suggesting an involvement of local adaptations in this region. For the SNP rs17868256, the derived G allele is associated with the Han Chinese trait of higher zygomatic arches and more cambered outwards and backwards zygomatics, and the corresponding allele frequency is also the highest in CHB (0.52), followed by UIG (0.36) and CEU (0.25), as indicates that rs 17868256 is involved in the phenotypic divergence in cheeks between Europeans and Han Chinese. The SNP rs61672954, associated with side face shapes was not replicated statistically in HAN-CZ (sample sizes for AA, GA genotypes were too small to analysis in UIG-R), but had the same facial variation patterns among different genotypes as in the discovery UIG-D cohort (Fig. S9). So we cannot remove their potential effect on facial shape.

One evident limitation of this study is the relative small sample sizes for the UIG cohorts. A power analysis revealed that tests would be constrained and the association signals would not be highly significant (e.g. all GWAS *P* values>1 × 10^−11^). And yet, even given the limited sample sizes we were still able to detect 6 genome-wide association signals, among which five were replicated to various degrees. This may be attributed mainly to the specific study designs: first, as the Uyghur was examined on the phenotypic dimensions where the ancestral groups EUR and HAN differ the most, the search was thus focused on genetic variants of large effect size, rendering higher test power for a given sample size. Second, the 3dDFM data densely annotates each face by over 30,000 vertices, resulting in virtually face phenome data. Based on this, the association signals were scanned both phenome-widely and genome-widely, as would greatly enhance the power of detecting the phenotype-genotype associations.

Several trends are notable involving the genetic architecture of facial morphology. First, no GWAS loci of “major effects” were identified that account for a large portion of phenotypic variance in spite of the strong overall divergence, e.g., in nasal shape across Eurasia ^2^ This is in contrast to the case of skin pigmentation whose major genetic factors explain substantial phenotypic variance ^42–44^. This suggests that the human face should be best described by a typical polygenic model of complex trait, characterized by a large number of variants of small effects. Second, most facial shape related variants seem to be pleiotropic. All the candidate loci in our study seem to be associated to the complex shape changes of whole face, not limited to the features of GWAS signals, in similar trends across the three sample cohorts, implying that such dispersive facial changes were induced by genetic variants rather than stochasticity (Figs. 3 and 4). At the individual gene level, *PCDH7* was also known for its versatile functions, related to not only mouth shape but also musical aptitude ^45^, waist-to-hip ratio ^46^ and many diseases ^40,41,47–49^ Another SNP rs3827760 in *EDAR,* replicated in this study was also known for its broad effects in hair morphology, incisor shape and sweat gland density ^26,27,50,51^. Third, the genetic effects of face related loci seem to be shared among different ethnicities ^52–54^. This is evident given that the association signals and facial patterns are in general consistent between Uyghur and Han Chinese (Tables 2 and 3, Fig. 4). It may be thus hypothesized that the stereotypic faces of ethnicities are merely result of population stratification of the face-related allele frequencies. For example, a European nose is “big” probably due to the co-segregation of a higher proportion of “big” nose alleles compared to that of an average Han Chinese. Indeed, four (rs118078182, rs60159418, rs61672954 and rs17868256) out of the six candidate SNPs in this study have moderate (*Fst_CEU-CHB_* > 0.08) to strong (for rs60159418, *Fst_CEU-CHB_* = 0.5, top 0.3% genome-widely) population differentiation, each of which seems to contribute a gradient to the continuous transition from European to Han Chinese faces.

In the end, we showed that an additive genetic model of whole face shape, based on a set of SNPs of top association signals, would lead to measurable predictive power. We showed that for independent individuals (UIG-R), the prediction model can also construct realistic 3D faces significantly closer to the actual face than random expectation (Figs. 5B and 5C); test in the hypothetic forensic scenarios revealed a moderate yet significant enhancement of the identification rate in males (Fig. 5D). The better performance in male face prediction than in females may attribute to the sexual dimorphism ^55,56^ and greater variation in male faces. If we measure the sample dispersion of individuals from the corresponding average faces by PSD, the dispersion was substantially larger in males (mean dispersion = 1.80) than in females (mean dispersion = 1.73, Kolmogorov-Smirnov test *P* = 0.0025). As on average the genetic distance from an individual to the average genome is equal for male and female (assuming only autosomal variants are concerned), the larger dispersion suggests on average bigger allelic effects in males than in females, thus better statistical power and prediction model in males. There may be redundant markers within the 277 top SNPs, as the SNP sets filtered for strong LD gave very similar performance (see Supplementary Note; Table S12 for prediction, Table S13 for forensic). Fundamentally, it can be argued that this prediction model is not really additive, as the best-fitting effect coefficient *α* is far lower than 1 (Fig. S7), even for the SNP set (209) well controlled for the physical LD (Fig. S7C). This may implicate epistatic interactions between the causal SNPs, or is probably mainly due to a proportion of false-positive signals in the top SNPs set. Further studies specifically designed for examination of the genetic architecture is needed to address this question. Furthermore, we noticed that increasing the number of top SNPs in the prediction model (either for *P* value < 1 × 10^−7^ or 1 × 10^−5^ SNPs) could not improve the predictive power. Such saturation analysis suggests a finite number of loci affecting the normal facial shape. After all, this prediction model is highly simplified and explorative. Much work is needed to improve its performance to be formally tested in real forensic scenarios.

## Materials and Methods

### Study cohorts

EUR was a resident cohort living in Shanghai with self-reported European ancestry between 16 and 42 years old. The HAN-TZ participants were self-reported Han Chinese samples collected from Taizhou, Jiangsu province. College students of self-reported Han ethnicity from Xiangnan University in Chenzhou, Hunan province were collected as HAN-CZ. The UIG-D and UIG-R were composed of college students of self-reported Uyghurs collected from Xinjiang Medical University in Urumchi, Xinjiang province. The self-reported ancestry information was requested for the last three generations, and individuals with mixed ancestry or missing information were excluded from further analyses. For EUR, a participant was used only if his/her ancestries of the last three generations were all from EU countries (as for 2015) plus Switzerland, Norway and Iceland. Individuals with obvious health problems or any history of facial surgery were ruled out. All sample collection in this study was carried out in accordance with the ethical standards of the ethics committee of the Shanghai Institutes for Biological Sciences (SIBS) and the Declaration of Helsinki, and has been specifically surveyed and approved by SIBS. All methods were carried out in accordance with the corresponding guidelines and regulations. A written statement of informed consent was obtained from every participant, with his/her authorizing signature. The participants, whose transformed facial images were used in this study as necessary illustrations of our methodology, have been shown the manuscript and corresponding figures. Aside from the informed consent for data sampling, a consent form stating the identifying information in an online open-access publication was shown and explained to each participant and their authorizing signature was obtained as well.

### High-density 3D facial images alignment

The 3dMDface^®^ system (www.3dmd.com/3dMDface) was used to collect high-resolution 3D facial images. We first established dense anatomical correspondence across dense surfaces of 3D facial images automatically as described previously ^12^. Briefly, 15 salient facial landmarks were annotated automatically based on the principal component analysis (PCA) projection of texture and shape information. A reference face was selected for high image quality and smooth surface, and its mesh was resampled to achieve an even density of one vertex in each 1mm × 1mm grid. There were 32,251 vertices in total for the reference mesh. Afterwards, the reference face was warped to register to each objective face to ensure the complete overlapping of the 15 landmarks via a thin-plate spline (TPS) transformation. The vertices on the reference face then found their closest projections on the sample face to define the samples' new vertices, resulting in a point-to-point correspondence. At last, the Generalized Procrustes analysis (GPA) was used to align the sample grids into a common coordinate system. As a result, we obtained a set of 32,251 3D points to represent each participant's face. Samples with defective images were removed from the study.

### Genotyping, quality control and imputation

Genomic DNA extracted from blood samples of UIG-D and UIG-R were genotyped on corresponding genotyping arrays. Quality control were performed using PLINK v1.07 ^57^. Furthermore, 92 individuals from UIG-D were whole-genome sequenced at high-coverage (30×). We didn't consider SNPs on mitochondria. SNPs with MAF <0.01, call rate <90%, or rejection in the Hardy-Weinberg Equilibrium test with *P*<1 × 10^−6^ were omitted from the study. Genomic ancestry was detected using EIGENSTRAT 5.0.2 ^58,59^ with CHB and CEU from 1000 Genomes Project ^60^ (1KG phase1 release v2) to remove samples who were not Uyghurs ancestry. Specifically, principal component analysis using 17,552 autosomal SNPs pruned from UIG-D panel based on call rate (>90%), MAF (>0.01), and LD (pairwise r2 <0.1) was used to assess population structure. Samples with genotype missing rate >0.1 were removed. Samples were further examined by pairwise IBD estimation and inbreeding coefficients to remove individuals of close genetic relationships. Specifically, we used 17,552 independent SNPs to estimate the pairwise IBD to find pairs of individuals who genetically look too similar to each other and set inbreeding coefficients >0.2 or <−0.2 as in inbreeding. Individuals with aberrant gender information checked using X chromosome data were discarded. Finally, samples with defective 3D images were removed from the study. Detailed results of QC were given in Table S5. In the end, a total of 847,046 SNPs were used in 694 UIG-D and 758,453 SNPs were used in the 171 UIG-R. Genomic DNA of HAN-CZ was extracted from saliva according to a modified Phenol-chloroform protocol ^61^. Targeted genotyping for the index SNPs were carried out by SNaPshot multiplex system on an ABI3130xl genetic analyzer by Genesky Biotech, Shanghai, China. The results of SNaPshot genotyping quality control were shown in Table S14.

In order to merge the data between UIG-D and UIG-R, which used different genotyping platform, we carried out imputation within UIG-D and UIG-R separately, using whole genome sequencing data as reference. Specifically, the chip data were pre-phased using SHAPEIT v2.r790 ^62^, and then imputed for probabilistic genotypes by Impute2 ^63^ taking 1,092 individuals from 1000 Genomes project phase 1 and 92 high-coverage (30×) whole-genome sequencing data of Uyghurs (unpublished data) as reference. The genotyped SNPs with a large difference between the info and concordance values (info_type0 – concord_type0 >0.1) or a low concordance value (<0.7) were excluded for further analysis. Additionally, SNPs were removed with low imputation quality score (<0.4), followed by converting the genotype probabilities to corresponding allele pair when it was over the threshold >0.9 using GTOOL software ^62^ SNPs with MAF <0.01 or a call rate <90% were omitted.

### Extraction of ancestry-divergent phenotypes

Three types of ancestry-divergent phenotypes were defined. The first type, the landmark based ancestry-divergent phenotypes were defined based on the 15 landmarks (Fig. S1A). Briefly, the whole face were divided into five parts (Table S1). Thirty six distances or angles were then defined as traditional anthropometric traits, many of which measure highly correlated facial features (Table S2). We chose 10 traits (Table S3) as the ancestry-divergent phenotypes, which surrogate the 36 traits and showed higher segregation (lower Students' test *P* value, Table S3) between EUR and HAN-TZ.

The second type is the PCA based ancestry-divergent phenotypes (sPC). Six facial features were extracted from the 3dDFM face data (Fig. 1B), which were then decomposed using PCA separately. The top PCs that together explained 98% total variance were examined for phenotypic segregation between EUR and HAN-TZ using Students' Test (Fig. S2 and Table S4). The PC that defined the most significant segregation was taken as the ancestry-divergent phenotype, named as sPC.

PLS based ancestry-divergent phenotypes (sPLS) were defined as follows: In the PLS equation “response ~ predictor”, we labeled the individuals from EUR as 1 and HAN-TZ as 2 in response. The predictor term featured 3 × n matrix of the 3dDFM data, where n stood for the number of vertices in the corresponding facial feature. The PLS regression were first carried out only with EUR and HAN-TZ data. An optimal PLS model is a regression model composed of the first *k* top ranking PLS component(s), where *k* renders the best prediction from predictor to response. Leave-one-out (LOO) was used to cross-validate PLS models, with increasing number of top components, and the root mean squared error of prediction (RMSEP) was used to evaluate the overall prediction. There exist two cross-validation (CV) estimates based on RMSEP (Fig. S3 and Table S4): ‘CV’ was the ordinary CV estimate, and ‘adjCV’ was a bias-corrected CV estimate. A PLS regression model achieves the maximal discrimination between EUR and HAN-TZ (Table S4) when the RMSEP showed turning point on adjCV curve (Fig. S3), and the corresponding *k* was used. An optimal PLS model was found in each facial feature and termed sPLS model.

### Statistical analyses and replications

Genome-wide association tests were carried out using PLINK v1.07 ^57^ and a linear regression model with additive SNP effects was used. For multiple testing correction, we performed the Benjamini-Hochberg procedure for FDR ^21^ to the combined results of the 66 autosomal GWA analyses. A FDR level of 0.05 was used to determine whether a test remained significant after multiple-testing correction.

For narrow-sense replication, the 3dDFM data from UIG-R and HAN-CZ were either projected to the sPC spaces or input to the sPLS models defined in UIG-D, to obtain the corresponding phenotype scores in narrow-sense replication. An additive linear regression was used to test the association between index SNPs and the corresponding narrow-sense phenotype. This test required a minimum of 10 individuals that carry at least one minor allele.

Index SNPs were also tested for broad-sense replication. For PLS-based permutation, genotypes (0, 1, 2 in additive model and 0, 1 in dominant model) of each SNP were taken as predictor. Principle components (PCs) that together explained 99% of the total variance of each facial feature were adopted as response. A scheme of Leave One Out (LOO) was used in which *N-1* individuals were used as training and the left-out was used as test. This was repeated until every individual was used as a test sample. The training set was used to build a PLS regression model whose optimal number of components was determined by RMSEP under cross-validation, as described before. Based on this model, a predicted genotype was given to the test sample. By such analogy, every sample obtained a predicted genotype. Afterwards, the correlation between actual genotype and predicted genotype can be calculated. Next, we performed a permutation procedure to control the supervised effect of PLS ^64–67^. Briefly, the genotypes were reshuffled randomly among the samples for 1,000 times, followed by the same procedure as above to calculate correlations between the actual and predicted genotypes. The null distribution of correlations can be thus established based on the permutation sets. The corrected PLS-based permutation *P* value was generated by ranking the empirical raw correlation against the null distribution.

For the pair-wise shape distance (PSD) permutation among genotypes ^13^, we calculated the Euclidean distances between the mean shapes of any two genotype groups. The mean shape can be denoted as a vector,

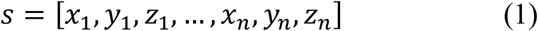

where *x_i_*, *x_i_*, *x_z_* was the X, Y, Z coordinate values of the *i*th points, *n* was the number of points.

For each two genotypes mean shape *s* and *s*′, the PSD was defined as

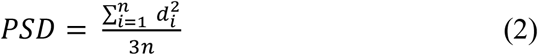

where 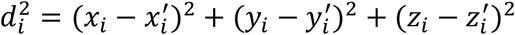.

We also randomly permuted the genotypes among the samples and then calculated the PSD between the peudo-genotype groups. The PSD scores resulted from permutation formed the null distribution. The one-side *P* value was calculated by the proportion of permuted PSD smaller than or equal to the true PSD.

### Data availability

All of the phenotypic measurements tested for association and 277 markers genotypes used for prediction are available through the Figshare, an online digital repository (https://figshare.com) with DOI: 10.6084/m9.figshare.4284770 (https://dx.doi.org/10.6084/m9.figshare.4284770).

## Acknowledgements

We are grateful to all the volunteers taking part in this study. We thank Chen Liu, Nianhao Cheng for the assistance in sample collection; thank Wei Qian for PLS-based Methods discussion; thank Bertram Müller-Myhsok, Benno Pütz for non-linear methods discussion; thank David Allen Hughes Jr. for manuscript polishing and comments.

## Author Contributions

Conceived and designed the study: K.T. Performed the study, summarized results and wrote the manuscript: L.Q., K.T. Advise the study and modified the manuscript: L.Q., S.W., S.X., K.T. Contributed reagents/material/software: Y.Y., P.F., S.H., H.Z., J.T., Y.L., H.L., D.L., S.W., J.G., S.P., L.J., Y.G., S.W., S.X. All authors read and approved the final manuscript.

